# The vasodilatory mechanism of nitric oxide and hydrogen sulfide in human mesenteric artery of colorectal cancer patients: role of potassium channels

**DOI:** 10.1101/666834

**Authors:** Awat Y. Hassan, Ismail M. Maulood, Abbas Salihi

## Abstract

Recent studies focused on the role of gasotransmitters in cancer progression and prevention. Therefore, this study was designed to explore the vasodilator activity of NO and H_2_S in human mesenteric artery of CRC patients via activation of K^+^ channels. For this purpose, two sets of experiments were established. The blood samples from CRC patients were obtained to detect serum levels of Endocan and MDA. Moreover, the role of K^+^ channels were assessed in mediating vasodilation of human mesenteric artery in response to SNP and Na_2_S. The level of serum Endocan was decreased in CRC patients compared to healthy individuals, while serum MDA was not changed. The arterial rings precontracted with NE were first relaxed by cumulative addition of increasing concentrations of either SNP (30nM-30μM) or Na_2_S (1-6mM). Then maximal relaxation rates were calculated for four times at each 15min intervals. Preincubation of arterial rings for 20min with individual K^+^ channels blockers were significantly reduced relaxation caused by SNP and Na_2_S at different time intervals. Furthermore, pretreatment of L-NAME did not change the vasodilation induced by Na_2_S. Vasodilation of CRC mesenteric unchanged by synergistic application of SNP and Na_2_S. While preincubation of arterial rings with PAG significantly enhanced vasodilation induced by SNP. In conclusion, these results indicate that endothelial dysfunction and oxidative stress do not take part in the pathogenesis of CRC. The dilatory mechanisms of NO and H_2_S in mesenteric arteries of CRC patients are K^+^ channels and time dependent, and the activity of CSE enzyme slows down the vasodilator ability of exogenous NO.

## Introduction

Colorectal cancer (CRC) is the third most frequently occurring malignancy and the second leading mortality cause of cancer worldwide. More than 1.8 million new CRC cases and 881,000 deaths was reported in 2018, accounting for about 1 in 10 cases and deaths [1]. Regardless of, the low incidence of CRC in the Middle East [2], the Kurdistan region of Iraq has a higher annual incidence rate of cancer, and it represents about 38 to 61.7 cases/100,000 population and it’s the fourth cancer in both sexes separately, and the cause of mortality of about 8.6% [3]. The most common histopathological type of CRC in this region is adenocarcinoma [4].

Endogenous gasotransmitters mainly nitric oxide (NO), carbon monoxide (CO), and hydrogen sulfide (H_2_S) are generated endogenously by specific enzymes [5]. NO is produced from L-arginine by the enzyme NO synthase (NOS) [6, 7] while H_2_S is synthesised from L-cysteine by either cystathionine beta-synthase (CBS), cystathionine gamma-lyase (CSE) or mercaptopyruvate sulfurtransferase (MST) [8, 9]. They can modulate different biological pathways and functions at a physiologically relevant concentration [10] by the opening of various kinds of membrane ion channels [11]. NO is an essential regulator of angiogenesis [12] and vasorelaxation through activation of guanylate cyclase and production of cyclic guanidine monophosphate (cGMP) [13]. cGMP, in turn, will increase an endothelial Ca^+2^ concentration and finally the opening of localised Ca^+2-^ dependent potassium (K_+_) channels (K_Ca_) [14]. In contrast, H_2_S exerts its effect on vasodilation [15] and angiogenesis [16] via direct activation of K_ATP_ channels [17].

Beside their independent actions, NO and H_2_S can act synergistically or antagonistically to elicit their downstream effects, ranging from their biosynthesis to the signalling cascade in a cell target. NO, and H_2_S are mutually dependent on regulating the vasodilation, angiogenesis [18, 19] and endothelial homeostasis [20] at different levels. The role of these gases in cancer is quite confusing, as both gases have tumour promotion and anti-tumour properties [21] and they can modulate various cancer cell functions, such as proliferation, invasion, metastasis, and tumour angiogenesis [22]. Moreover, the enzymes responsible for NO and H_2_S production are upregulated in CRC cells, endogenously produced low-to-mid concentrations of H_2_S or NO support cell proliferation, while exogenous delivery of H_2_S or NO can suppress the colon cancer cells division [23].

Intra-tumoral blood vessels are of paramount for tumour growth, metastasis and therapy [24] and the acquisition of a differential reactivity by functional, mature blood vessels in the tumour microenvironment represents a proper target for antitumor therapy [25]. Because cancer cells are exhibiting an accelerated metabolism, therefore they are demanding high reactive oxygen species (ROS) concentrations to maintain their high proliferation rate [26]. Thus, the high levels of ROS can damage or kill cells by oxidising proteins, lipids, and nucleic acids [27]. Overall, these observations elucidate that oxidative stress and cancer are closely linked [28]. A growing number of researches is investigating the consequences of ROS and endothelial dysfunction in cancer [29]. Nonetheless, to the best of our knowledge, no researcher has linked between oxidative stress, endothelial function and vascular reactivity to NO and H_2_S in CRC patients. Therefore, the study aimed to evaluate endocan as an endothelial function marker and MDA as an oxidative stress marker in CRC patients. Also we were aimed to evaluate the mechanisms responsible for NO and H_2_S induced vasodilation in human mesenteric artery of CRC patients. To approach, these questions, possible roles of different K^+^ channels in the vasodilation response produced by NO and H_2_S has been assessed.

## Subjects and Methods

### Subjects

This study is a case-control study. The study was approved by the human ethics committee belonging to College of Science, Salahaddin University-Erbil, and the informed consent from all the patients was obtained. For the first experiment, the subjects were recruited in two hospitals in Erbil city, Rizgray hospital Oncology department and Nanakaly Hospital. Venous blood was taken from 44 CRC patients with different stages of colorectal cancer (24 males and 20 females). Also, 40 healthy volunteers of similar age were recruited in this study as healthy individuals. While for the second experiment colorectal tumour specimen were obtained from colorectal cancer patients undergoing partial colectomy at CMC and welfare private hospitals in Erbil city.

Patients were selected according to histological evidence of colonic or rectal adenocarcinoma when ages of patients greater than 18 years and they were able to donate blood. The blood samples were obtained by phlebotomy under aseptic technique. Blood was put into clot activator tube for serum separation, and then the sera were separated under 2000rpm centrifugation for 5min. In contrast, patients with underlying immunodeficiency disorder or immunodeficiency state and individuals who had other co-morbid health problems which could introduce heterogeneity to the sample, (e.g. arthritis, asthma, diabetes mellitus, hypertension, and other inflammatory diseases) were excluded.

## Methods

### Determination of Endocan

The concentration of Endocan was determined according to Sandwich-ELISA method. The micro ELISA plate was pre-coated with an antibody specific to Endocan. Standards or samples were added to the suitable micro ELISA plate wells and combined with the specific antibody. Then a biotinylated detection antibody specific for Endocan and Avidin-Horseradish Peroxidase conjugate and substrate were added to each microplate well. Only those wells that contain Endocan, biotinylated detection antibody and Avidin-HRP conjugate appeared blue in colour and reaction was stopped with 1N H_2_SO_4_ solution, and the colour turns yellow. The absorbance was measured spectrophotometrically at a wavelength of 450nm.

### Determination of Serum Malondialdehyde

Malondialdehyde (MDA) was determined according to Ohkawa method [30]. The procedure was started by TBA preparation; in which 0.66g of Thiobarbituric acid (TBA) dissolved in 100 ml of 0.05M of NaOH with simple heating. Then, Trichloroacetic acetic acid (TCA) was prepared by dissolving 17.5g of TCA in 100 ml distilled water; while TCA2 was prepared by dissolving 70 g of TCA in 100 ml distilled water. Finally, 150 ml of serum was added 1ml of TCA1 mixed for 2 min and incubated in boiling water bath for 15 min, then 1 ml of TCA2 was added and incubated for 20 min at 37C° then centrifuged for 5 min at 2000 rpm the supernatant was read at 532 nm.

## Myographical recording

### Vessel Collection and preparation

Human mesenteric arteries from patients suffering from CRC and underwent partial colectomy was collected. The arteries supplying blood to the tumour were dissected out surgically and directly were added to a beaker containing cold modified Krebs solution (Fig 1) and aerated with 95% O_2_ and 5% CO_2_. In the laboratory, the excess tissue and fat were removed, and the arteries were cut into rings about 3-4 mm long.

**Fig 1.**
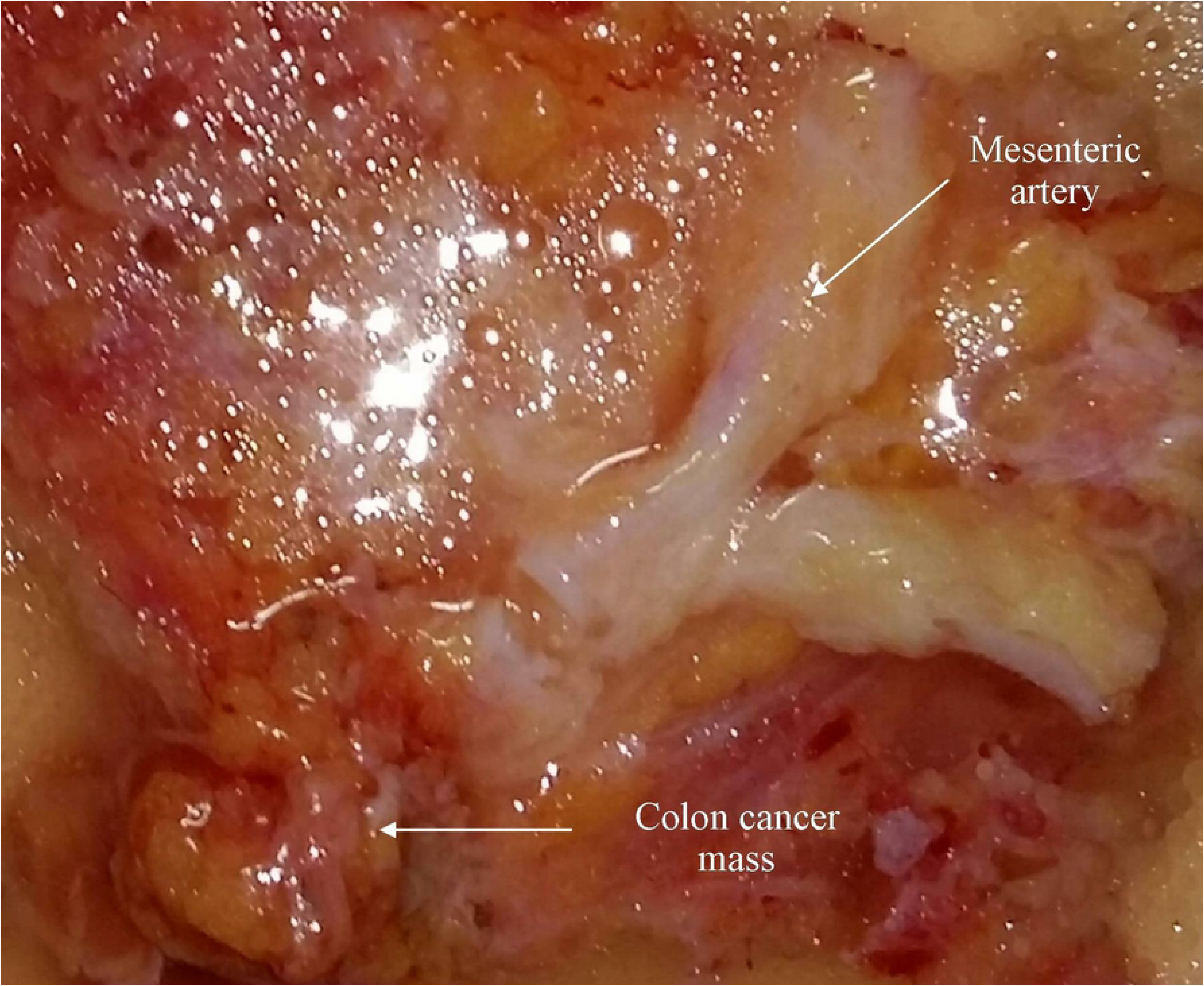
Mesenteric artery feeding colon tumour.

### Recording of Isometric tension

The procedure of [31] with some modifications was followed to study the vasodilator activity of the isolated mesenteric arteries. The arterial rings were held up by two stainless steel clamps. The first clamp was attached to a hook at the underside of the organ bath jacket and the other was connected to the force transducer through a thread to record the isometric tension of the mesenteric arteries, and the data was recorded by LabChart data acquisition software. The held up arterial rings were immersed in Krebs solution (NaCl 5.10gm/L, NaHCO_3_ 1.94/L, MgSO_4_ 0.686gm/L, KCl 2.24gm/L, KH_2_PO4 0.15gm/L, CaCl_2_ 0.277gm/L, C6H12O6 2gm /L), contained in a 10 ml organ chamber. Krebs solution was maintained at pH 7.4 and was constantly aerated with 95% O_2_ 5% CO_2_ at 37°C (Panlab Harvard Apparatus, USA).

The mesenteric arterial rings were tensed to a stable basal strain of 4gm before left to equilibrate for 2hrs. To remove cellular metabolites, Krebs solution was replaced every 15-20min intervals in the bath chamber until it reaches the stability and the experimental substances acquaint with the bath chambers according to the protocols, the arteries were incubated with drugs for 20 minutes then norepinephrine (NE) 1μM until reaches maximum contractility and plateau, then relaxation happened by bolus single dose application of sodium nitroprusside (SNP) or sodium disulfide (Na_2_S).

### Experimental protocol

The arterial rings precontracted with NE were first relaxed by cumulative addition of either SNP (30nM-30μM) or Na_2_S (1-6 mM). Based on these initial experiments, relative half-inhibitory concentration (IC_50_) of SNP (2.3μM), Na_2_S (2.4mM) had retested for the ability to relax precontracted rings in three separate sets of experiments. In the first experiment, when the NE-induced contraction reached the uppermost value, SNP (2.3μM) or Na_2_S (2.4mM) was added and left for 60min, and the maximal relaxation rate (%) was calculated four times at each 15min intervals (n=8). Then the role of K^+^ channels in the progress of SNP and Na_2_S mediated relaxation were tested via incubation of the arterial rings for 20min with the Tetraethylammonium (TEA;1mM), Glibenclamide (GLIB;0.1μM), Barium Chloride (BaCl_2_; 1mM) and 4-aminopyridine (4-AP;1mM). While, in the second experiment, the role of endogenous NO and H_2_S were tested by preincubation of arterial rings with either L-NAME (3×10^-4^M) or D,L-propargylglycine (PAG) (10mM), antagonists of eNOS and CSE for 20min before applying SNP (n=8). Finally, to examine whether the combination of H_2_S and NO potentiate or inhibit the vasorelaxation when the NE-induced contraction reached the highest value, SNP and Na_2_S were added simultaneously and left for 60 min, (n=8).

### Statistical analysis

The statistical analysis of myographical data was performed using two-way analysis of variance (ANOVA) followed by Bonferroni *post hoc* test. Maximum relaxation responses were calculated as a percentage of the contraction produced by NE and were expressed as the means±standard error of the mean (SEM). The tension created by NE was defined as 0% relaxation, and the baseline tension before the addition of NE was determined as 100% relaxation. While, comparison between CRC and healthy individuals were made by Mann-Whitney test and the values were represented as median and quartiles. A p-value less than 0.05 (P<0.05) were considered statistically significant. The graphs, calculation and statistical analysesp were performed using GraphPad Prism software 6.0 (GraphPad Software, San Diego, California, USA).

## Results

### Serum Endocan and Malondialdehyde concentration

Serum endocan concentration was markedly lower in the CRC patients (67.56, 43.04-94.28) than the healthy individuals (88.68, 59-101.3). While, there were no significant differences in MDA concentration between the CRC patients (3.62, 2.86-4.96) and the healthy individuals (3.78, 3.23-4.24), Fig 2A and B, respectively.

**Fig 2.**
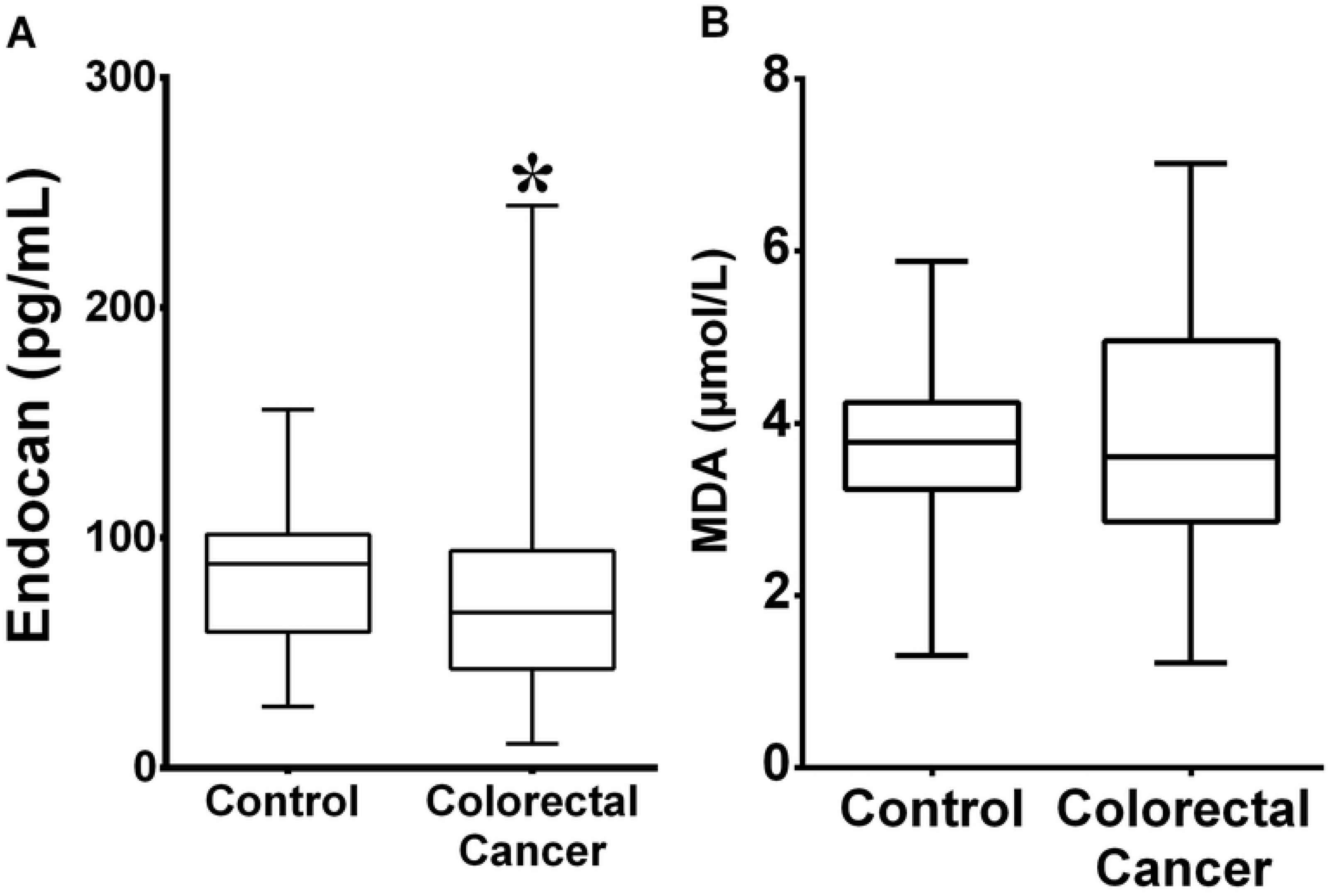
Comparison between Endocan (pg/mL), MDA (μmol/L) level in a healthy group and CRC patients. Endocan significantly (*P<0.05,) decreased in CRC patients compared to the healthy control group, while Non-significant change was observed in MDA level. The comparison was performed using the unpaired Mann-Whitney test.

### Measurement of IC_50_

Sodium nitroprusside at concentrations from 30nM-30μM caused a relaxant effect on mesenteric arteries of CRC when precontracted with NE (1 μM) with IC_50_’s of 2.42±0.16 μM (IC_50_ of CI 95% between 1.18 to 4.95 μM), and the percentage of relaxation was 80.74±7.256%. While Na_2_S at concentrations from 1-6 mM had a relaxant effect on mesenteric arteries of CRC precontracted with NE, the calculated IC_50_’s was 3.54±1.07mM (IC_50_ of CI 95% between 1.4 to 5.68mM), and the percentage of relaxation was 84.43±22.05%. Concentration-response curve for the effect of SNP and Na_2_S against NE-mediated contractions are shown in Fig 3A and B, respectively.

**Fig 3.**
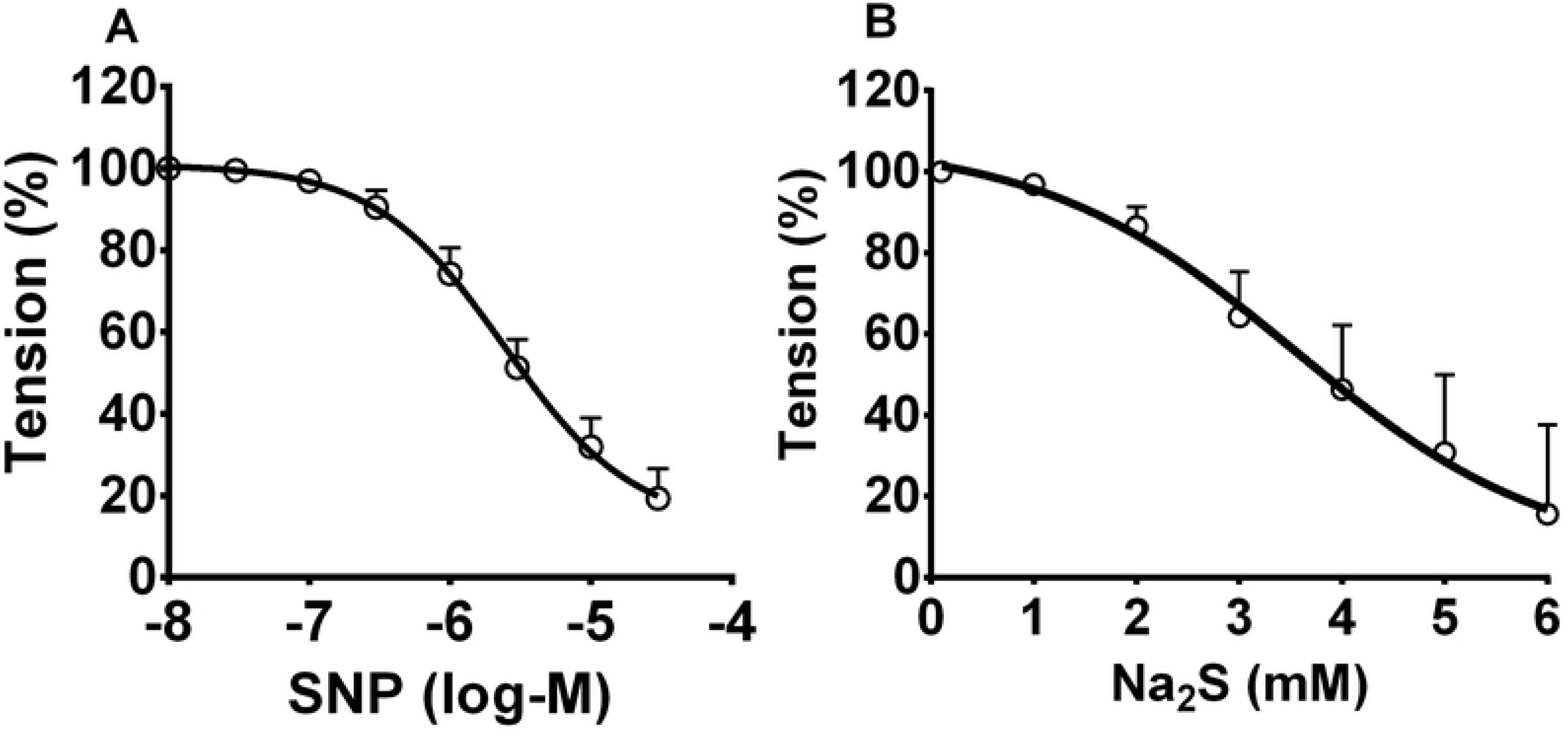
Cumulative dose-response curve for the vasorelaxant effects of (A) SNP (30nM-30μM); and (B) Na_2_S (1-6mM) on NE (1μM) induced contraction in mesenteric arteries of CRC patients.

### The role of K^+^ channels in the NO-induced relaxation

To identify the role K^+^ channels on the time-dependent change of CRC feeding mesenteric arteries relaxation responses to effect of SNP, arterial rings were incubated with K^+^ channels blockers for 20 min before the addition of SNP and the net vasorelaxation effect of SNP in mesenteric arteries was measured every 15min over the course of 60 min. Preincubation of mesenteric arteries with either GLIB (10μM, n=6), TEA (1 mM, n=6) or 4-AP (1 mM, n=6) remarkably reduced the net vasorelaxation effect of SNP in mesenteric arteries at all-time intervals. While, the reduction of vasorelaxation responses of SNP was lasted for 45 min after preincubation of the mesenteric arterial ring with BaCl_2_ (1mM, n=6), as shown in Table 1 and Fig 4.

**Fig 4.**
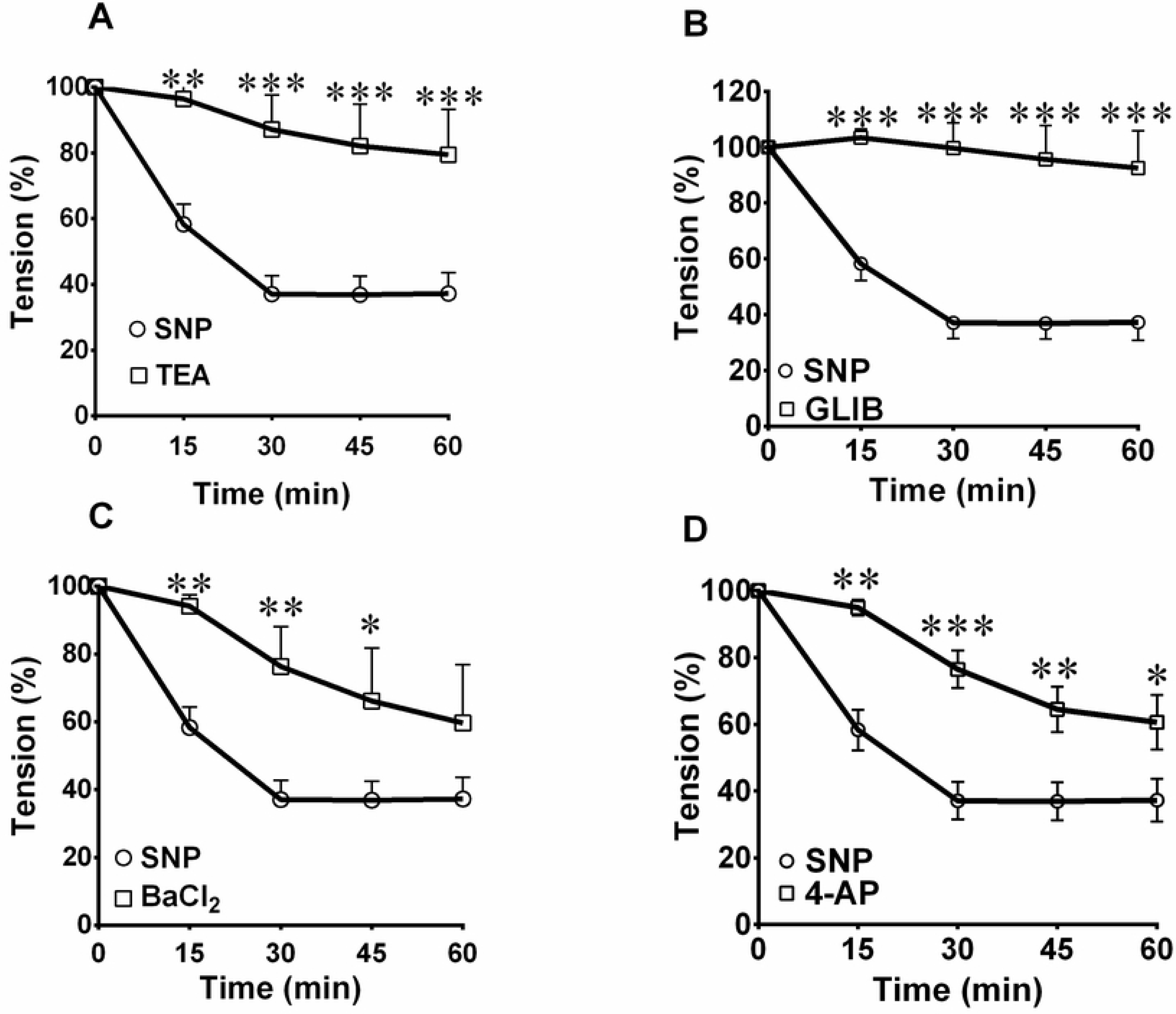
Time-dependent change of relaxation responses to effect of SNP in mesenteric arteries preincubated with TEA (1mM), BaCl_2_ (1mM), 4-AP (1mM), GLIB (10μM) respectively. (A) SNP-induced vasorelaxation significantly inhibited by TEA (▪) pretreatment (**P<0.01, 5 min, *** P<0.001, 30, 45, 60min). (B) SNP-induced vasorelaxation significantly inhibited by GLIB (▪) pretreatment (*** P<0.001, 15, 30, 45,60min). (C) SNP-induced vasorelaxation significantly inhibited by BaCl_2_ (▪) pretreatment (**P<0.01, 15 and 30min, (*P<0.05, 45 min). (D) SNP-induced vasorelaxation significantly inhibited by 4-AP (▪) pretreatment (**P<0.01, 15 and 45 min, **P<0.01, 30min, *P<0.05, 60 min).

**Table 1.** Involvement of K^+^ channels in the mechanism of time-dependent relaxation responses to effect of SNP in mesenteric arteries of CRC patients.

|  | Time |  |  |  |
| --- | --- | --- | --- | --- |
|  | 15 min | 30 min | 45 min | 60 min |
| **Control** | 41.7±6.09 | 62.9±5.64 | 63.1±5.65 | 62.7±6.38 |
| **TEA** | 3.6±4.52 | 12.9±10.5 | 17.9±12.7 | 20.6±13.8 |
| **GLIB** | 0.1±3.14 | 0.3±9.04 | 4.4±12.2 | 7.4±13.4 |
| **BaCl<sub>2</sub>** | 5.9±3.39 | 23.7±11.8 | 33.9±15.6 | 40.4±17.2 |
| **4-AP** | 5.1±2.39 | 23.5±5.67 | 35.5±6.77 | 39.4±8.15 |

### The role of K^+^ channels in the H_2_S-induced relaxation

The impairment of Na_2_S induced mesenteric artery relaxation taken from CRC patients was continued throughout the experiment when the arteries were previously incubated with either GLIB (n=6) or 4-AP (n=6). While BaCl_2_ reduced vasorelaxation responses produced by Na_2_S only in the middle of the experiment. In contrast, TEA had failed to amend the vasorelaxation responses of Na_2_S, as shown in Table 2 and Fig 5.

**Fig 5.**
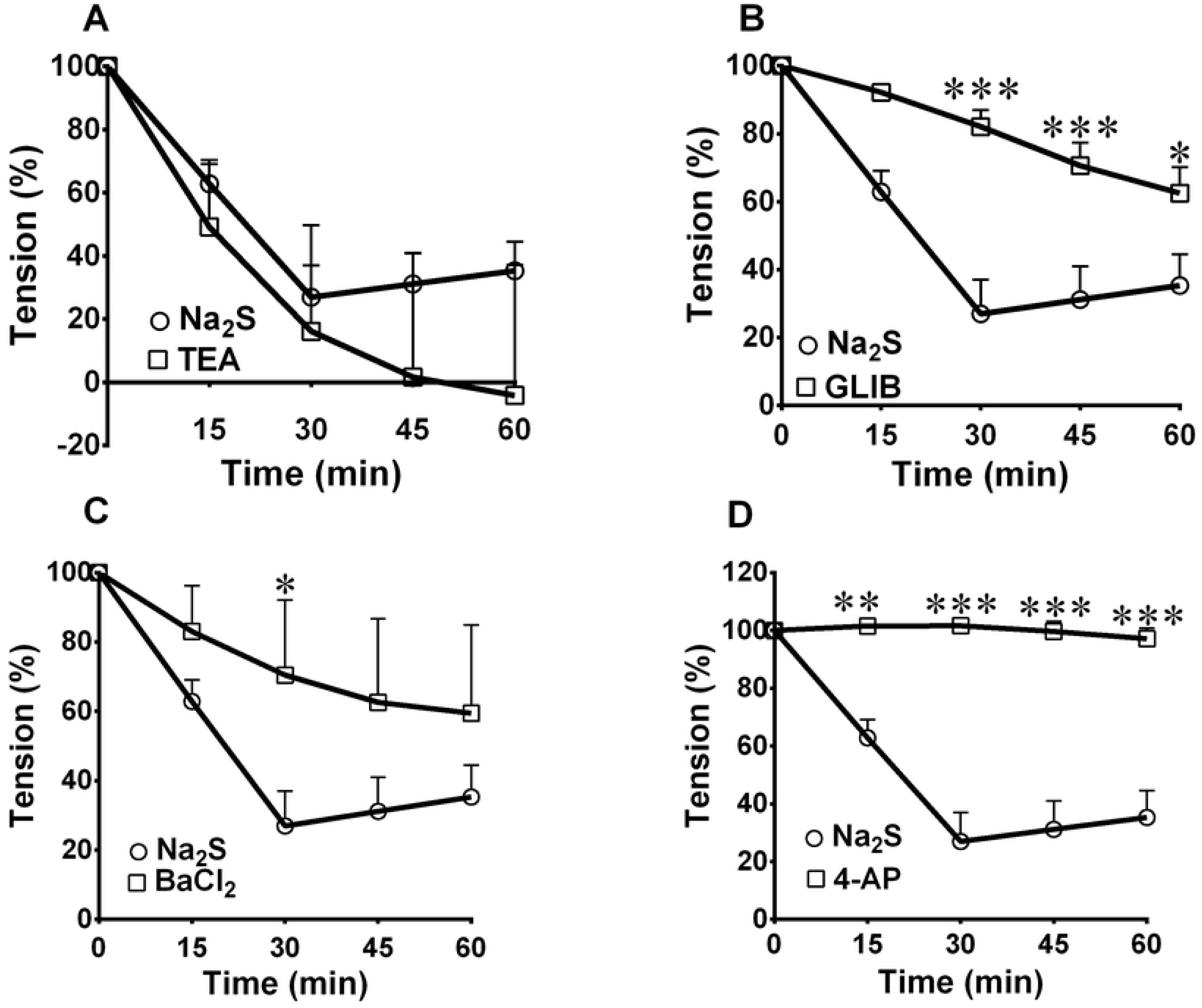
Time-dependent change of relaxation responses to the effect of Na_2_S in mesenteric arteries preincubated with TEA (1mM), GLIB (10μM.), BaCl_2_ (1mM), 4-AP (1mM) respectively. (A) TEA (▪) had no significant effect on Na_2_S-induced time-dependent arterial relaxation. (B) Na_2_S-induced vasorelaxation significantly inhibited by GLIB (▪) pretreatment (*** P<0.001, 30, 45, min, * P<0.05, 60min). (C) Na_2_S - induced vasorelaxation significantly inhibited by BaCl_2_ (▪) pretreatment (* P<0.05, 30min). (D) Na_2_S-induced vasorelaxation significantly inhibited by 4-AP (▪) pretreatment (** P<0.01, 15min, *** P<0.001, 30, 45, 60min).

**Table 2.** Involvement of K^+^ channels in the mechanism of time-dependent relaxation responses to effect of Na_2_S in mesenteric arteries of CRC patients.

|  | Time |  |  |  |
| --- | --- | --- | --- | --- |
|  | 15 min | 30 min | 45 min | 60 min |
| **Control** | 37.1±6.26 | 73±10.1 | 68.8±9.85 | 64.7±9.21 |
| **TEA** | 13.9±4.52 | 81.3±10.5 | 105±12.7 | 116±13.8 |
| **GLIB** | 7.8±1.91 | 17.9±4.82 | 29.4±6.71 | 37.5±7.61 |
| **BaCl<sub>2</sub>** | 17±13.2 | 29.5±21.7 | 37.4±24.1 | 40.5±25.4 |
| **4-AP** | 2±1.34 | 2±2.6 | -0.3±3.53 | 3.9±3.72 |

### Interaction effects of SNP and Na2S

Combination of SNP and Na_2_S did not modify relaxation responses at all time-intervals in comparison to the relaxation induced by individual application of either SNP or Na_2_S. Furthermore, preincubation of arterial rings with L-NAME (n=6) did not change the extent of the Na_2_S-induced relaxation at all time-courses of the study. On the other hand, treating the mesenteric arterial rings with PAG (n=6) significantly increased vasorelaxation induced by SNP at all time-courses of the study, as shown in Table 3 and Fig 6.

**Fig 6.**
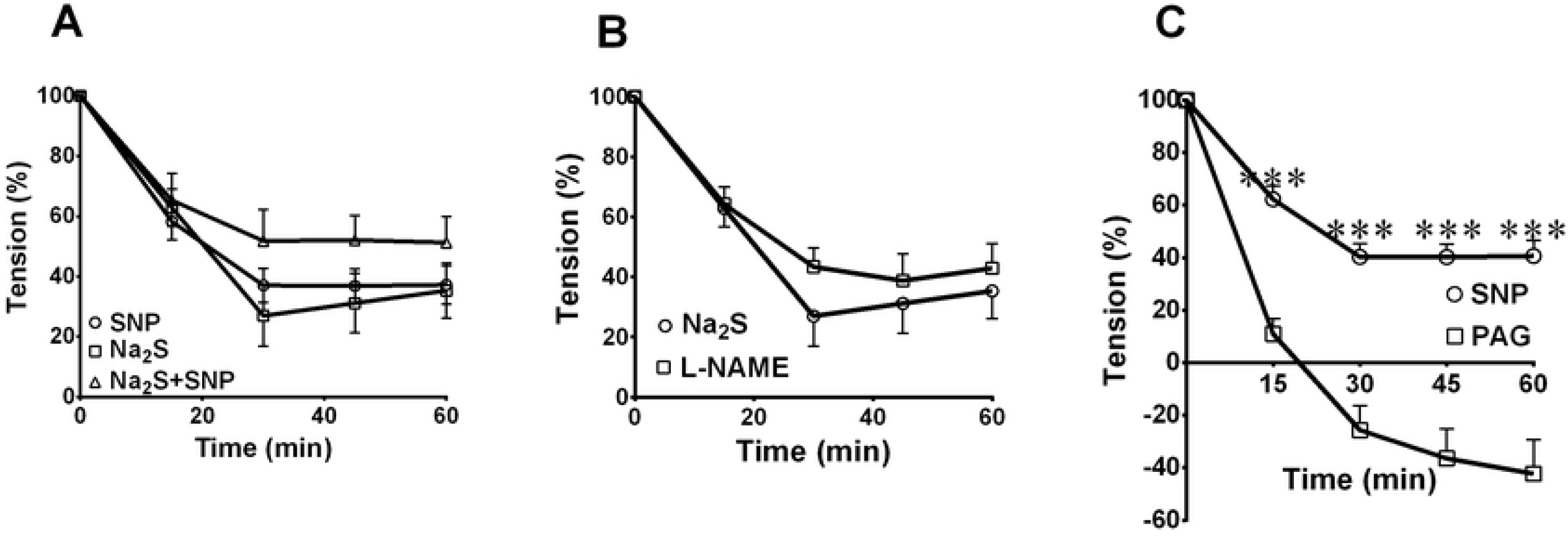
The combination effects of SNP and Na_2_S, L-NAME and PAG preincubation on the time-dependent relaxation responses of the mesenteric arteries precontracted with NE. (A) The combination of SNP and Na_2_S (▲) did not change the time-dependent relaxation. (B) L-NAME (0.3mM; ▪) had no significant effect on Na_2_S-induced time-dependent relaxation. (C) SNP-induced relaxation significantly inhibited by PAG (10mM; ▪) pretreatment (*** P<0.001, 15, 30,45,60min).

**Table 3.** The combination effects of SNP and Na_2_S, L-NAME and PAG preincubation on the time-dependent relaxation responses of the mesenteric arteries of CRC patients.

|  | Time |  |  |  |
| --- | --- | --- | --- | --- |
|  | 15 min | 30 min | 45 min | 60 min |
| **SNP+Na<sub>2</sub>S** | 34.7±8.96 | 48.2±10.4 | 47.9±8.09 | 48.7±8.67 |
| **L-NAME** | 35.8±5.73 | 56.8±6.23 | 61.2±8.87 | 57.1±8.3 |
| **PAG** | 89±5.87 | -25.7±9.42 | -36.5±11.3 | -42.3±12.9 |

## Discussion

The first part of the present study ascertains that endothelium cells of CRC patients are functioning normally, because the serum Endocan level markedly decreased, which is an endothelial cell marker [32]. The reason of endocan decrement was explained by [33–35], that is due to either chemotherapy or VEGF receptor-2 kinase inhibitors treatment or the downregulation of endocan expression, suggesting that the expression of endocan is related to the development and differentiation of CRC [32]. While, [36] stated that Endocan correlates with colon tumour size, depth of invasion, lymph node metastasis, distant metastasis and Dukes’ staging. Also, there has been a growing interest in studying MDA as a marker of oxidative stress in cancer progression [37]. In the present, we showed that the serum MDA level was not changed in CRC patients comparing to healthy individuals. Although, previous studies had been observed that MDA is increased in CRC patients [38], but [39] explained that chemotherapeutical treatment would normalize the oxidative stress in CRC patients. Taken together, these results indicate that CRC patients have intact arteries and their endothelium are functioning normally.

In the second part of the present study, we found that SNP profoundly relaxes mesenteric arteries in CRC patients. In this concern, the previous study conducted by [40] recorded 69% relaxation of rat mesenteric arteries, whilst the relaxation of human hand veins reached 103% [41]. Then, to explore the exact mechanism of relaxation of SNP, the role of K^+^ channels were tested in the mesenteric arteries of CRC patients.

The results of the present study unveil that K^+^ channels have a great role in the SNP-induced relaxation because pretreatment of mesenteric arteries with either TEA, GLIB, BaCl_2_ or 4-AP markedly delayed vasodilation. NO activates several K^+^ channels of SMCs in rat, rabbit mesenteric artery and cerebral arteries, including ATP-sensitive K^+^ channels (Deka and Brading 2004; Koh et al. 1995; Murphy and Brayden 1995) and induce membrane hyperpolarization by decreases in [Ca^2+^]_i_ levels through inhibition of Ca^2+^ influx or Ca^2+^ release from intracellular stores [42]. Moreover, NO hyperpolarize arterial smooth muscle cells via activation both K_V_ and K_Ca_ channels on VSMC in rat superior mesenteric artery, coronary and cerebral artery and large artery [43–45], through a cGMP-dependent mechanism, in that way inhibiting the evoked membrane depolarization and upsurge in [Ca^2+^]_i_ [46]. Furthermore, this result is in accordance with the earlier finding of Schubert, Krien (47), they were observed that VSMC K_IR_ currents are regulated by NO. Meanwhile, Hempelmann, Seebeck (48) displayed that neither 4-AP nor BaCl_2_ modulate NO-induced relaxation in rat basilar artery. This result proposes that NO might exert vasodilation possibly by the opening of different K^+^ channels. Hence, we can conclude that K^+^ channels play a central role in the vasodilation mechanism of NO.

Consistently, we were the first to observe that Na_2_S potently relax mesenteric arteries in CRC patients. Recently, Materazzi, Zagli (49) exhibited that H_2_S could relax precontracted human mesenteric arterial rings in a concentration-dependent manner. Moreover, we found that the relaxation of mesenteric arteries induced by Na_2_S depends on the activation of K_ATP_ and K_V_ channels. The importance of K_ATP_ channel activation confirms in a human colonic mesenteric artery what has already been observed in the rat arterial smooth muscle by [50] human mammary artery by [51], either through hyperpolarization of SMCs membrane, in turn meight close voltage-gated Ca^2+^channels [52] or through channel protein sulfhydration [15]. In the same way, in the rat aorta H_2_S induces vasorelaxation was diminished by KCNQ-type K_V_ channels blockage [53]. In contrast [54] had found that K_ATP_ channels did not mediate the relaxations caused by H_2_S in the guinea-pig ileum and trout urinary bladder. Even though, K_IR_ channels weakly participated in the relaxation of mesenteric arteries in CRC patients, while previous studies conducted by [55–57] were reported that the mechanism of relaxation in rat aorta is mainly mediated by the stimulation of K_IR_ channels and subsequent K_IR_-dependent hyperpolarization from the endothelium to the smooth muscle cells

Although, H_2_S was found to activate BK_Ca_ [58], IK_Ca_, and SK_Ca_ channels in endothelial cells [15]; and BK_Ca_ channels in SMCs of mesenteric arteries [58] and cerebral arterioles [59]. The results of the present study disclosed that TEA did not change the vasodilation of mesenteric arteries induced by Na_2_S, signifying that K_Ca_ might not be accountable for H_2_S induced vasorelaxation. More or less the same results were observed by [60], who noted that different blockers for K_Ca_ channels ineffective in the vascular effect of H_2_S. Opposite to our results [55] found that the maximum relaxation of VSMC of rat artery [15, 61] and human mammary artery [51] induced by NaHS was significantly attenuated by K_Ca_ channels blockers.

Because both H_2_S and NO are vasorelaxant factors, and they have dissimilar mechanisms of action, one may predict an improver effect when the two gases are applied together [50], whereas the results of the current study showed that the combination of SNP and Na_2_S donors were not changed maximal relaxation during all time course of the study compared to SNP and Na_2_S respectively. One explanation is a combination of H_2_S, and NO generate a new molecule (perhaps nitrosothiol), which does not relax blood vessels either *in vitro* or *in vivo* [62]. Consequently, the formation of this novel molecule most likely denotes a means for biological inactivation or perhaps sequestration of released NO [63]. In contrast, rat aortic relaxation was prolonged when both gas donors are added together [64]. This synergistic action may be due to the production of HSNO and HNO as a result of a chemical reaction between H_2_S and nitrite [65], which releases NO and polysulfides and relax VSMCs through soluble guanylyl cyclase activation [66].

At the same time, preincubation of L-NAME did not change the relaxation effects induced by Na_2_S. Similar results have been reported by Ohia, Opere (67); Monjok, Kulkarni (68) they were notified that L-NAME did not modify the relaxation effects of Na_2_S in isolated porcine irides, this means that endogenous NO has no impact on the vasoactivity of H_2_S donor. In contrast, preincubation of CRC arterial rings with PAG increased relaxant activity induced by NO donor. The proper explanation for this result is that endogenous H_2_S inhibits the action of NO. In the same manner, SNP-induced vasorelaxation of rat aorta and human internal mammary artery was diminished by low concentration of H_2_S by suppressing NO action or inhibition of NO synthase [52, 60].

## Conclusion

In conclusion, low Endocan and normal MDA values reveal that endothelial dysfunction and oxidative stress do not take part in the pathogenesis of CRC. Beside this, the mechanism of NO and H_2_S-induced mesenteric artery vasodilation is time and K^+^ channels dependent, in which NO dilates mesenteric arteries via activation of K_ATP_, K_Ca_, K_IR_ and K_V_ channels, while vasodilation activity of H_2_S is due to the modulation of K_ATP_ and K_V_ channels. Moreover, NO and H_2_S are interacting at the level of enzymes, the activity of CSE enzyme slows down the vasodilator ability of exogenous NO.

## Acknowledgements

Our special thanks are due to all staff members of the Oncology department at Rizgary and Nanakaly hospital for their help, and our highest appreciation to Dr Saeb Gailany and Dr Imad at CMC and Welfare hospitals for providing the human patient colonic specimens.

